# The MASLD-causing E167K variant of TM6SF2 is degraded by the ubiquitin ligase RNF5

**DOI:** 10.64898/2026.10.21.760790

**Authors:** Emad Heidary Arash, Huimin Pan, Holger Willenbring

## Abstract

The TM6SF2 E167K loss-of-function variant is one of the most penetrant genetic risk factors for metabolic dysfunction-associated steatotic liver disease (MASLD), including progression to cirrhosis and hepatocellular carcinoma. Here we show that TM6SF2 E167K is recognized and degraded by the ER-associated degradation pathway, and identify RNF5 as the ubiquitin ligase tagging TM6SF2 E167K, providing a rational target for MASLD therapy.

## Introduction

Metabolic dysfunction-associated steatotic liver disease (MASLD) is a major health challenge because it is common and complicated by severe hepatic fibrosis (cirrhosis) and hepatocellular carcinoma (HCC)^1^. Genome-wide association studies have identified genetic risk factors that predispose to MASLD, with a variant in transmembrane 6 superfamily member 2 (*TM6SF2*) converting glutamic acid 167 to lysine (E167K) being particularly pathogenic^2,3^. E167K has been shown to cause TM6SF2 protein degradation^2^, potentially because of misfolding^2,4^, but how TM6SF2 E167K is degraded—the molecular root of its pathogenicity—is unknown.

The clinical phenotypes caused by the TM6SF2 E167K variant are replicated by TM6SF2 deficiency, indicating loss of function^5^. In accord, exogenous hepatic expression of wild-type (WT) TM6SF2 and, to a lesser extent, the E167K variant reduces steatosis, fibrosis and HCC in *Tm6sf2*-deficient mice^5^. Therefore, identification of the molecular mechanism underlying TM6SF2 E167K degradation would provide a rational target for therapy of MASLD in patients carrying this variant.

TM6SF2 is an endoplasmic reticulum (ER) membrane protein known to regulate lipid secretion through very low-density lipoprotein (VLDL) particles^4^. Newly synthesized proteins entering the secretory pathway are subject to ER-associated degradation (ERAD), a quality control pathway that eliminates misfolded proteins through the ubiquitin-proteasome system^6^. Because of the physical separation of misfolded ER-localized proteins from the cytoplasmic proteasome complex, ERAD substrates need to be extracted through the ER membrane in a process referred to as retrotranslocation, which relies on the hexameric AAA ATPase complex p97/VCP^6^.

## Results

Confirming previous findings^2^, we found that TM6SF2 E167K affords lower protein levels than TM6SF2 WT at similar mRNA levels (Fig. 1 A and B). To determine whether TM6SF2 E167K is degraded by the ubiquitin-proteasome system or the autophagy-lysosomal pathway^7^, we treated FLAG-TM6SF2-expressing HEK293T cells with the proteasome inhibitor MG132 or Bafilomycin A, which blocks lysosomal protein degradation. While MG132 stabilized TM6SF2 E167K to levels comparable to TM6SF2 WT, Bafilomycin A had no effect (Fig. 1 C and D). Neither drug changed TM6SF2 mRNA expression (Fig. 1E). In accord, immunoprecipitation of FLAG-TM6SF2 co-expressed with HA-Ubiquitin showed that TM6SF2 E167K was much more ubiquitinated than TM6SF2 WT (Fig. 1F). These findings show that the lysine substitution at residue 167 in TM6SF2 results in misfolding and degradation by the ubiquitin-proteasome system.

**Fig. 1.**
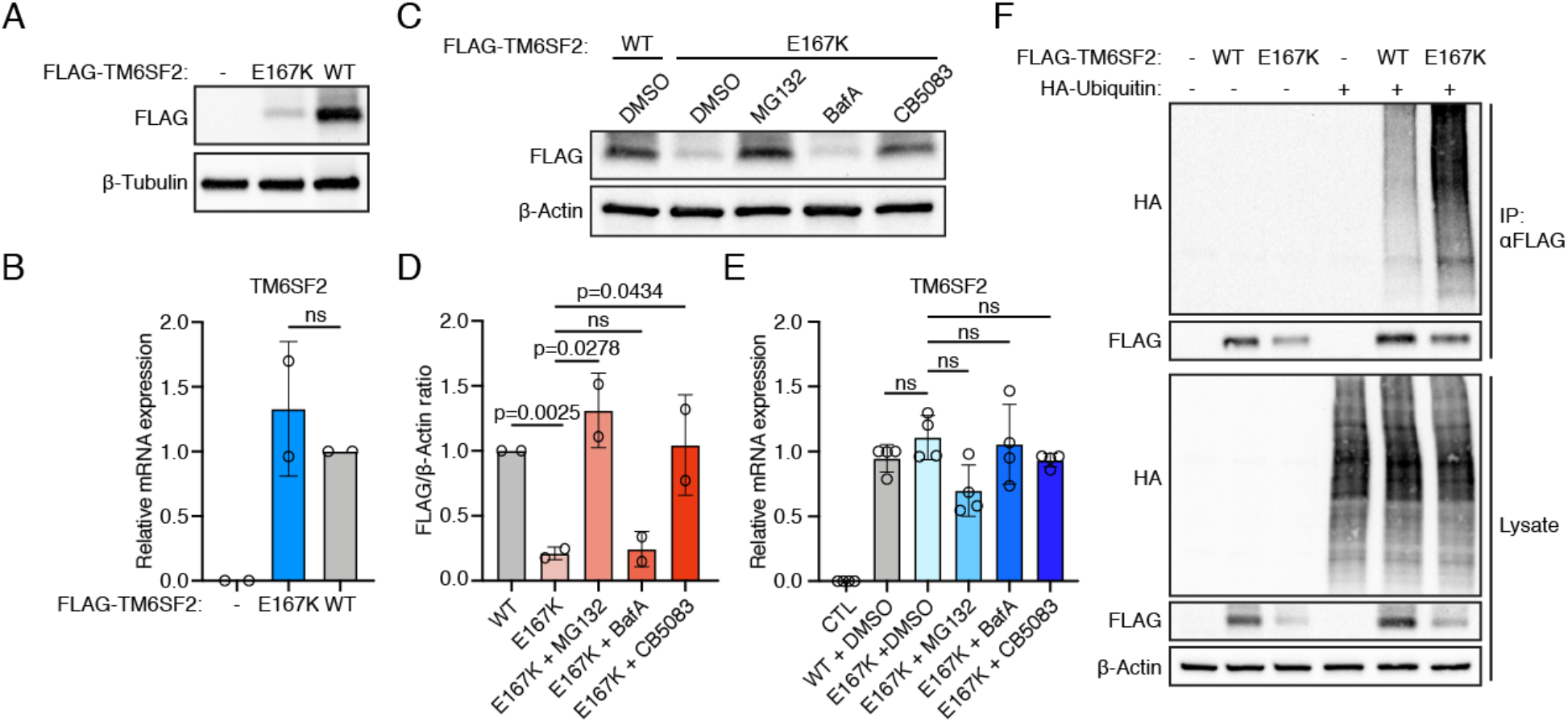
TM6SF2 E167K is a misfolded protein degraded by the ubiquitin-proteasome. (A and B) Immunoblotting (A) and qRT-PCR (B) of HEK293T cells transfected with FLAG-TM6SF2 WT or FLAG-TM6SF2 E167K (mean ± SD, n = 2 independent experiments, one-way ANOVA). (C-E) Immunoblotting (C) with quantification of FLAG to β-Actin ratio (D) and qRT-PCR (E) of HEK293T cells transfected with FLAG-TM6SF2 WT or FLAG-TM6SF2 E167K and treated with 10 μM MG132, 100 nM Bafilomycin A (BafA) or 5 μM CB5083 (mean ± SD, n = 2 independent experiments, one-way ANOVA). (F) Immunoblotting of HEK293T cells co-transfected with FLAG-TM6SF2 WT or FLAG-TM6SF2 E167K and HA-Ubiquitin and immunoprecipitated with anti-FLAG antibody.

To determine whether TM6SF2 E167K is an ERAD substrate requiring p97/VCP-mediated retrotranslocation, we treated FLAG-TM6SF2 E167K-expressing HEK293T cells with the p97/VCP inhibitor CB5083, which stabilized the protein without changing mRNA expression (Fig. 1 C-E). Immunoprecipitation of FLAG-TM6SF2 E167K showed binding to p97/VCP, which was enhanced by CB5083, indicating that TM6SF2 E167K and p97/VCP are trapped in a complex within the membrane when retrotranslocation is blocked (Fig. 2A). These findings show that TM6SF2 E167K is an ERAD substrate and retrotranslocated by the p97/VCP complex for proteasomal degradation.

**Fig. 2.**
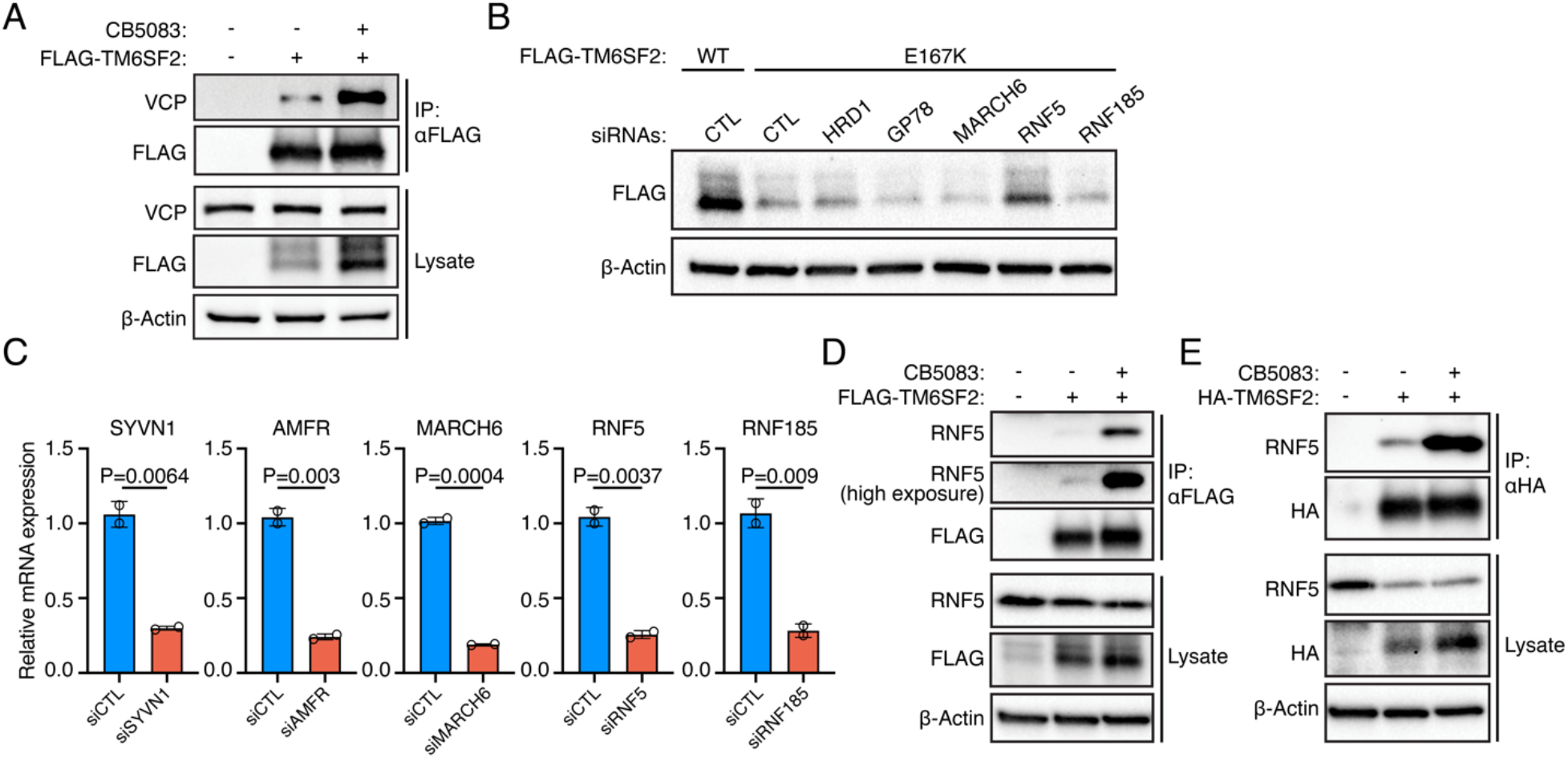
TM6SF2 E167K is an ERAD substrate targeted by the ubiquitin ligase RNF5. (A) Immunoblotting of HEK293T cells transfected with FLAG-TM6SF2 E167K and treated with DMSO or 5 μM CB5083 and immunoprecipitated with anti-FLAG antibody. (B and C) Immunoblotting (B) and qRT-PCR (C) of Huh7 cells co-transfected with FLAG-TM6SF2 WT or FLAG-TM6SF2 E167K and the indicated siRNAs targeting ERAD ubiquitin ligases (mean ± SD, n = 2 biological replicates, two-tailed t-test). (D and E) Immunoblotting of HEK293T cells transfected with FLAG-TM6SF2 E167K (D) or HA-TM6SF2 E167K (E) and treated with DMSO or 5 μM CB5083 and immunoprecipitated with anti-FLAG or anti-HA antibodies.

To identify the ubiquitin ligase(s) that degrade TM6SF2 E167K, we performed a loss-of-function screen of ERAD ubiquitin ligases known to target integral membrane proteins^6^. We performed the screen in Huh7 cells to ascertain liver-disease relevance. Although HRD1 (encoded by *SYVN1*) and its closely related homolog GP78 (encoded by *AMFR*) collectively degrade all luminal and multiple transmembrane ER proteins and are therefore the most important ubiquitin ligases of the ERAD pathway^6,8^, siRNA-mediated depletion of either ligase did not increase TM6SF2 E167K protein levels (Fig. 2 B and C). Depletion of MARCH6, which targets multiple misfolded membrane proteins^6,8^, including disease-causing NPC1 variants^9^, also had no effect. Only depletion of RNF5 prevented TM6SF2 E167K protein degradation. Depletion of the closely related ubiquitin ligase RNF185 had no effect, which is consistent with previous findings that RNF5 and RNF185 target different ERAD membrane substrates despite sequence similarity^10^. To confirm the role of RNF5 in recognition of TM6SF2 E167K, we immunoprecipitated FLAG-and HA-tagged TM6SF2 E167K, which showed binding to endogenous RNF5 (Fig. 2 D and E). This physical interaction was enhanced by CB5083, indicating that TM6SF2 E167K remains trapped in a complex with RNF5 when retrotranslocation is inhibited.

## Discussion

Here we find that the MASLD-causing E167K variant of TM6SF2 encodes an unstable and polyubiquitinated protein that is recognized by the ERAD pathway for degradation and identify RNF5 as the ubiquitin ligase that tags TM6SF2 E167K for degradation. RNF5 is known to tag the F508del variant of the cystic fibrosis transmembrane conductance regulator (CFTR) for degradation^11^, suggesting RNF5 inhibition as a therapeutic strategy for cystic fibrosis^12^. In accord, a common ancestral haplotype associated with lower expression of RNF5 has been shown to be protective against bacterial colonization in patients with cystic fibrosis^13^. Thus, our findings suggest siRNA-mediated depletion, which proved to be effective and safe in recent clinical trials for other diseases of hepatic lipid metabolism^14,15^, or pharmacological inhibition of RNF5 as a therapy for MASLD in patients carrying the TM6SF2 E167K variant.

## Materials and methods

### Antibodies and reagents

The following primary antibodies were used for immunoblotting at the indicated concentrations: Mouse monoclonal FLAG-M2 (Sigma-Aldrich, F3165; 1:2000), rabbit monoclonal HA-tag (Cell Signaling Technology, 3742; 1:2000), mouse monoclonal RNF5 (22B3) (Santa Cruz Biotechnology, sc-81716; 1:1000), rabbit polyclonal VCP (Cell Signaling Technology, 2648; 1:1000), rabbit monoclonal β-Actin (13E5) (Cell Signaling Technology, 4970; 1:5000), rabbit polyclonal β-Tubulin (Cell Signaling Technology, 2146; 1:5000). The following small molecules were used: MG132 (Selleck Chemicals, S2619), Bafilomycin A (Tocris Bioscience, 1334), CB5083 (MedChemExpress, HY-12861).

### Cell lysis, immunoblotting and immunoprecipitation

Cells were washed with PBS and lysed with ice-cold lysis buffer (50 mM Tris, pH 7.4, 150 mM NaCl, 1% NP40 and Halt protease inhibitor cocktail (Thermo Fisher Scientific, 78429). Lysates were cleared by centrifugation at 20,000 x g for 20 minutes at 4°C and 4x Laemmli sample buffer containing 10% β-mercaptoethanol was added to the supernatant. For immunoprecipitation, anti-FLAG or anti-HA antibodies were added to the cleared lysates and samples were rotated overnight at 4°C. Magnetic Dynabeads Protein G (Invitrogen, 10003D) was added to lysates and incubated for 1 hour with rotation at 4°C and washed three times with washing buffer (50 mM Tris-HCl pH 7.4, 150 mM NaCl, 0.1% Triton X-100) before adding 2x Laemmli sample buffer containing 10% β-mercaptoethanol. Cell lysates or immunoprecipitated proteins were resolved by SDS-PAGE on Mini-PROTEAN TGX Precast Protein Gels (Bio-Rad) and transferred to PVDF (Sigma-Aldrich, IPVH00010) or nitrocellulose (Bio-Rad, 1620112) membranes, and blocked with 5% nonfat dry milk in TBST (Tris-buffered saline supplemented with 0.1% Tween-20) at room temperature for 1 hour. Membranes were incubated with primary antibodies diluted in blocking buffer (5% nonfat dry milk or BSA in TBST) overnight at 4°C and washed three times with TBST before incubation with HRP-conjugated antibodies diluted 1:5000 in blocking buffer at room temperature for 1 hour. After washing three times with TBST, membranes were imaged using a Bio-Rad ChemiDoc imaging system after adding ECL substrate (Thermo Fisher Scientific).

### Cell culture and transfection

HEK293T cells were purchased from ATCC (CRL-3216) and cultured in DMEM supplemented with 10% FBS and antibiotics. Huh7 cells were obtained from the Japanese Collection of Research Bioresources Cell Bank (JCRB0403) and cultured in RPMI 1640 supplemented with 5% FBS and antibiotics. Cells were maintained at 37°C in 20% O2 and 5% CO2. Plasmids were transfected with Lipofectamine 3000 (Thermo Fisher Scientific, L3000001) according to the manufacturer’s instructions. siRNAs were transfected with either Lipofectamine RNAiMAX (Thermo Fisher Scientific, 13778150) or Lipofectamine 3000 according to the manufacturer’s instructions.

### RNA interference

To silence gene expression, siRNAs were transfected as a pool of oligonucleotides at a final concentration of 20 nM. The following siRNAs were purchased from Dharmacon: siCTL (siGENOME D-001206-14-05); siMARCH6 (siGENOME D-006925); siAMFR (siGENOME D-006522); siRNF5 (siGENOME D-006558); siSYVN1 (siGENOME D-007090). RNF185 siRNAs were purchased as dicer-substrate siRNA (DsiRNA) duplex oligonucleotides from IDT.

### RNA extraction and quantitative real time PCR

RNA was extracted using PureLink RNA Mini Kit (Thermo Fisher Scientific, 12183025) according to the manufacturer’s instructions. RNA was reverse transcribed to cDNA using qScript cDNA SuperMix (Quantabio, 95046) and qRT-PCR was performed on an Applied Biosystems ViiA 7 machine with SYBR Green master mix (Thermo Fisher Scientific, 4309155) using the following primers: SYVN1: Fwd: 5’GCTCACGCCTACTACCTCAAA3’, Rev: 5’GCCAGACAAGTCTCTGTGACG3’; AMFR: Fwd: 5’TTCGTCGGCACAAGAACTATC3’, Rev: 5’GCACAGTCGTCATTGTTGACAG3’; RNF5: Fwd: 5’GCCAGAACGGCAAGAGTGT3’, Rev: 5’GGCTCCCTCGCCCATAAAG3’; RNF185: Fwd: 5’GTGTTTACATCAGTGGTTGGAGA3’, Rev: 5’GTGCTGCCCCTTCCATAGAG3’; MARCH6: Fwd: 5’CACCTGAGAAACCGCTTTATCA3’, Rev: 5’CCGTGAAGGCATATCTGGAGA3’; GAPDH: Fwd: 5’TGCACCACCAACTGCTTAGC3’, Rev: 5’GGCATGGACTGTGGTCATGAG3’. Relative mRNA expression was determined by the ΔΔCt method with normalization to GAPDH.

### Statistical analysis

Data were analyzed using Prism GraphPad software and *P* values were determined using one-way ANOVA or Student’s t-test as indicated in figure legends.

## Acknowledgements

The authors thank Jacquelyn Maher for discussion and manuscript editing. This work was supported by NIH RC2 DK136052 and P30 DK026743 (UCSF Liver Center) to H.W. and an NIH T32 DK60414 postdoctoral fellowship a UCSF IRM Postdoc Independent Research Grant to E.H.A.

## Author contributions

E.H.A. conceived the project, performed experiments, analyzed and interpreted the data; H.P. performed experiments; H.W conceived and supervised the project, and interpreted the data;

E.H.A. and H.W. wrote the manuscript.

## Competing interests

The authors declare no competing interests.

## Notes

### Competing Interest Statement

The authors have declared no competing interest.

## References

1. Targher, G., Valenti, L., and Byrne, C.D. (2025). Metabolic Dysfunction-Associated Steatotic Liver Disease. N Engl J Med 393, 683–698.

2. Kozlitina, J., Smagris, E., Stender, S., Nordestgaard, B.G., Zhou, H.H., Tybjærg-Hansen, A., Vogt, T.F., Hobbs, H.H., and Cohen, J.C. (2014). Exome-wide association study identifies a TM6SF2 variant that confers susceptibility to nonalcoholic fatty liver disease. Nat Genet 46, 352–356.

3. Liu, Y.L., Reeves, H.L., Burt, A.D., Tiniakos, D., McPherson, S., Leathart, J.B., Allison, M.E., Alexander, G.J., Piguet, A.C., Anty, R., et al. (2014). TM6SF2 rs58542926 influences hepatic fibrosis progression in patients with non-alcoholic fatty liver disease. Nat Commun 5, 4309.

4. Luo, F., Oldoni, F., and Das, A. (2022). TM6SF2: A Novel Genetic Player in Nonalcoholic Fatty Liver and Cardiovascular Disease. Hepatol Commun 6, 448–460.

5. Newberry, E.P., Hall, Z., Xie, Y., Molitor, E.A., Bayguinov, P.O., Strout, G.W., Fitzpatrick, J.A.J., Brunt, E.M., Griffin, J.L., and Davidson, N.O. (2021). Liver-Specific Deletion of Mouse Tm6sf2 Promotes Steatosis, Fibrosis, and Hepatocellular Cancer. Hepatology 74, 1203–1219.

6. Christianson, J.C., Jarosch, E., and Sommer, T. (2023). Mechanisms of substrate processing during ER-associated protein degradation. Nat Rev Mol Cell Biol 24, 777–796.

7. Pohl, C., and Dikic, I. (2019). Cellular quality control by the ubiquitin-proteasome system and autophagy. Science 366, 818–822.

8. Leto, D.E., Morgens, D.W., Zhang, L., Walczak, C.P., Elias, J.E., Bassik, M.C., and Kopito, R.R. (2019). Genome-wide CRISPR Analysis Identifies Substrate-Specific Conjugation Modules in ER-Associated Degradation. Mol Cell 73, 377–389 e311.

9. Schultz, M.L., Krus, K.L., Kaushik, S., Dang, D., Chopra, R., Qi, L., Shakkottai, V.G., Cuervo, A.M., and Lieberman, A.P. (2018). Coordinate regulation of mutant NPC1 degradation by selective ER autophagy and MARCH6-dependent ERAD. Nat Commun 9, 3671.

10. van de Weijer, M.L., Krshnan, L., Liberatori, S., Guerrero, E.N., Robson-Tull, J., Hahn, L., Lebbink, R.J., Wiertz, E., Fischer, R., Ebner, D., and Carvalho, P. (2020). Quality Control of ER Membrane Proteins by the RNF185/Membralin Ubiquitin Ligase Complex. Mol Cell 79, 768–781 e767.

11. Younger, J.M., Chen, L., Ren, H.Y., Rosser, M.F., Turnbull, E.L., Fan, C.Y., Patterson, C., and Cyr, D.M. (2006). Sequential quality-control checkpoints triage misfolded cystic fibrosis transmembrane conductance regulator. Cell 126, 571–582.

12. Sondo, E., Falchi, F., Caci, E., Ferrera, L., Giacomini, E., Pesce, E., Tomati, V., Mandrup Bertozzi, S., Goldoni, L., Armirotti, A., et al. (2018). Pharmacological Inhibition of the Ubiquitin Ligase RNF5 Rescues F508del-CFTR in Cystic Fibrosis Airway Epithelia. Cell Chem Biol 25, 891–905 e898.

13. D’Antonio, M., Reyna, J., Jakubosky, D., Donovan, M.K., Bonder, M.J., Matsui, H., Stegle, O., Nariai, N., D’Antonio-Chronowska, A., and Frazer, K.A. (2019). Systematic genetic analysis of the MHC region reveals mechanistic underpinnings of HLA type associations with disease. Elife 8.

14. Nissen, S.E., Ni, W., Shen, X., Wang, Q., Navar, A.M., Nicholls, S.J., Wolski, K., Michael, L., Haupt, A., Krege, J.H., and Investigators, A.T. (2025). Lepodisiran - A Long-Duration Small Interfering RNA Targeting Lipoprotein(a). N Engl J Med 392, 1673–1683.

15. Fabbrini, E., Rady, B., Koshkina, A., Jeon, J.Y., Ayyar, V.S., Gargano, C., DiProspero, N., Wendel, S., Hegge, J., Hamilton, H., et al. (2024). Phase 1 Trials of PNPLA3 siRNA in I148M Homozygous Patients with MAFLD. N Engl J Med 391, 475–476.

